# Confined environment facilitates stacked conformations in Holliday Junction

**DOI:** 10.1101/2023.06.17.545413

**Authors:** Priya Agarwal, Sahil Hasan Kabir, Nibedita Pal

## Abstract

Holliday Junction (HJ) is an important intermediate for homologous recombination and is involved in DNA break repair. It is a highly dynamic structure and transitions between two stacked isomers through an intermediate open conformer. In a constrained cellular environment, HJ is expected to behave differently than in vitro. This can affect target recognition by junction-binding proteins and their subsequent functions. However, the studies on the effect of the confined environment on HJ is extremely scanty. In this work, by employing fluorescence-based techniques we investigated the effect of confinement on HJ conformation using reverse micelle encapsulation as a model for confined space. We observed that inside reverse micelle, HJ prefers to adopt stacked conformation. Most strikingly, even at lower divalent cation concentrations, the preference for stacked conformation is prevalent over an open conformation. Our finding suggests that such confinement-induced changes in the conformer population might affect the interaction and activity of the HJ-recognizing proteins in the cellular environment.

## INTRODUCTION

Four-way Holliday Junction (HJ) is involved in important cellular processes such as homologous recombination and DNA break repair.^1,2^ The very dynamic nature of HJ allows it to adopt two broad classes of conformers, an open X-like structure and a stacked structure where two arms are coaxially stacked on each other (Fig. 1(A)).^3–5^ The open X-like structure predominantly exists at low salt concentrations due to the electrostatic repulsion between the arms. HJ with non-homologous sequences adopts two isomers and *iso-I* and *iso-II* which can be differentiated from different stacking arms (Fig. 1(A)). At higher salt concentrations, HJ transitions between these two isomers through the intermediate open conformation (Fig. 1(A)).^5,6^ While the isomerization rate largely depends on the salt (mainly divalent cation) concentration, the relative population of the isomers is solely determined by the sequence at the junction, and it is practically impossible to selectively populate one of the isomers for a particular junction sequence.^7^

**Fig. 1.**
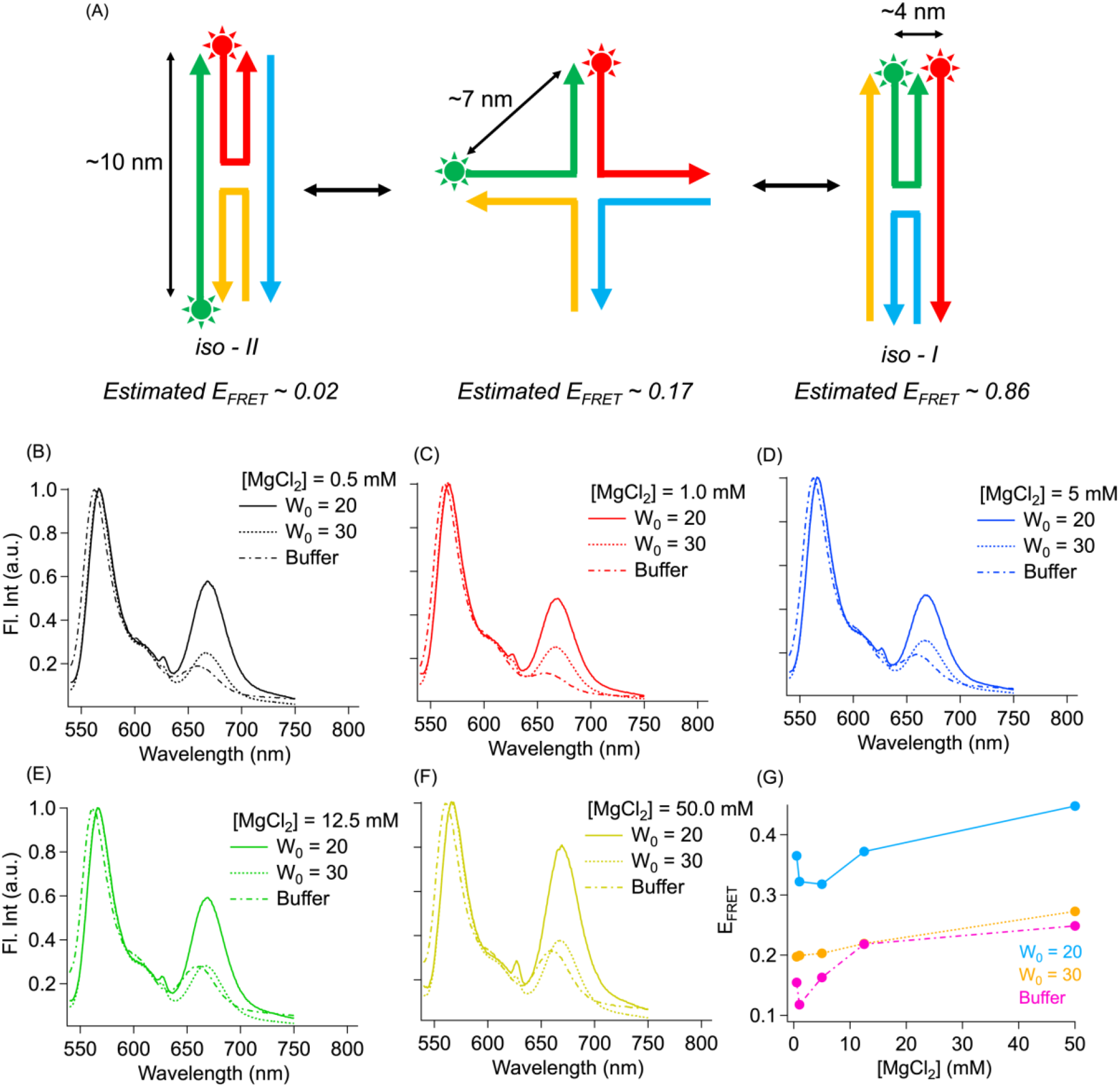
(A) Fluorophore labeling scheme of the J1. The Cy3-Cy5 distance are approximately calculated from dsDNA base-pair distances. Steadystate fluorescence spectrum of Cy3-Cy5 labeled J1 in buffer and encapsulated inside RMs with (B) 0.5 mM, (C) 1.0 mM, (D) 5 mM, (E) 12.5 mM and (F) 50 mM MgCl_2_. (G) Change in FRET efficiency with MgCl_2_ in different environment.

HJ relies on its dynamic nature to carry out its function which often involves proteins and enzymes binding to the junction. Junction-binding proteins and enzymes are found to bind both the stacked isomers in sequence independent-way. It is observed that enzymes bind to the stacked conformers as a dimer but unfold/distort them to a different extent to carry out their biological functions.^4,8^ On the other hand, the open X-like structure is recognized by tetrameric motor proteins.^9^ More interestingly, X-like structure recognizing proteins are generally found to be sequence-specific whereas proteins that recognize stacked conformers are not.^9^

Although single-molecule fluorescence techniques are extensively used to characterize the isomerization dynamics of HJ, the experiments are mostly carried out in aqueous buffer solution.^4–6,8^ On the contrary, the cellular environment is highly confined/restricted due to the high content of other macromolecules. Thus, the knowledge of how HJs behave in a restricted environment is scanty.^10^ This information is of paramount importance as the prevalence of one conformer over another (if any) can impact junction recognition by proteins and enzymes and thus the kinetic rate of downstream processes.

In this work, we have aimed to address this lacuna and explored the effect of confinement on HJ conformers. We have selected HJ J1, (see Materials and Methods for sequence) which has been extensively used in 3D self-assembly of DNA nanostructures.^11–13^ This junction (J1) has also been investigated in aqueous buffer by single-molecule and ensemble fluorescence experiments.^7,14^ These studies showed that at lower MgCl_2_ concentrations (2-5 mM) J1 preferentially populates *iso-I*. At much higher MgCl_2_ concentrations, however, both *iso-I* (60-70%) and *iso-II* (30-40%) are populated with a bias towards *iso-I*. It is noteworthy that these experiments were carried out in dilute aqueous buffer conditions and the effect of restricted/confined environment has not been investigated. As confinement effect influences both the structural and dynamic organization of the biomolecules, deviating from that in dilute solution,^10,15,16^ it will be important to know if this junction shows a bias towards any conformers in such an environment. Restricted environments are generally created by adding artificial crowding agents, such as (poly)ethylene glycol, dextran etc.^10,15,16^ However, reverse micelles (RM) offer an excellent confined environment without the need for adding any artificial agent to create confinement. RM carries a polar water pool inside lined by an amphiphilic layer of surfactants. and has been extensively used to study the confinement effect on biomolecules.^17–19^ The water-pool size can be varied in a systematic way by changing amounts of water, thus varying the amount of confinement. In this study, we used sodium bis(2-ethylhexyl)sulfosuccinate (AOT) RM to investigate the effect of confined environment on HJ. We have employed fluorescence-based techniques and observed that in a confined environment, J1 prefers stacked conformation, even at a lower MgCl2 concentration. This is contrary to the findings at dilute conditions as at lower salt concentrations HJ is known to adopt an open X-like structure. Additionally, our study reveals that in a confined environment structural heterogeneity of HJ is minimized compared to that in aqueous buffer. We anticipate that observation might have an important biological implication in HJ recognition process by proteins and enzymes in a confined cellular environment.

## MATERIALS & METHODS

### HJ assembly

HPLC purified fluorescently labeled and unlabeled oligonucleotides (sequences in Table 1) were purchased from Integrated DNA Technologies and Eurofins Scientific and were used without purification. 2-8 µM stock of J1 was prepared by mixing the stands at 1:1:1:1 ratio in 20 mM Tris-HCl (Sigma-Aldrich), pH 7.5 with an requied concentration of MgCl_2_. Two types of HJs were annealed, one with both donor (Cy3) and acceptor (Cy5) fluorophores containing S1, S2, S3, and S4 and another with only donor containing S1, S3, S4, and S5. HJs were annealed by mixing the four strands by heating the mixture to 90°C for 15 minutes and slowly cooling it down to 25°C for ∼5h, followed by incubation at 4°C till further use.

**Table 1:**
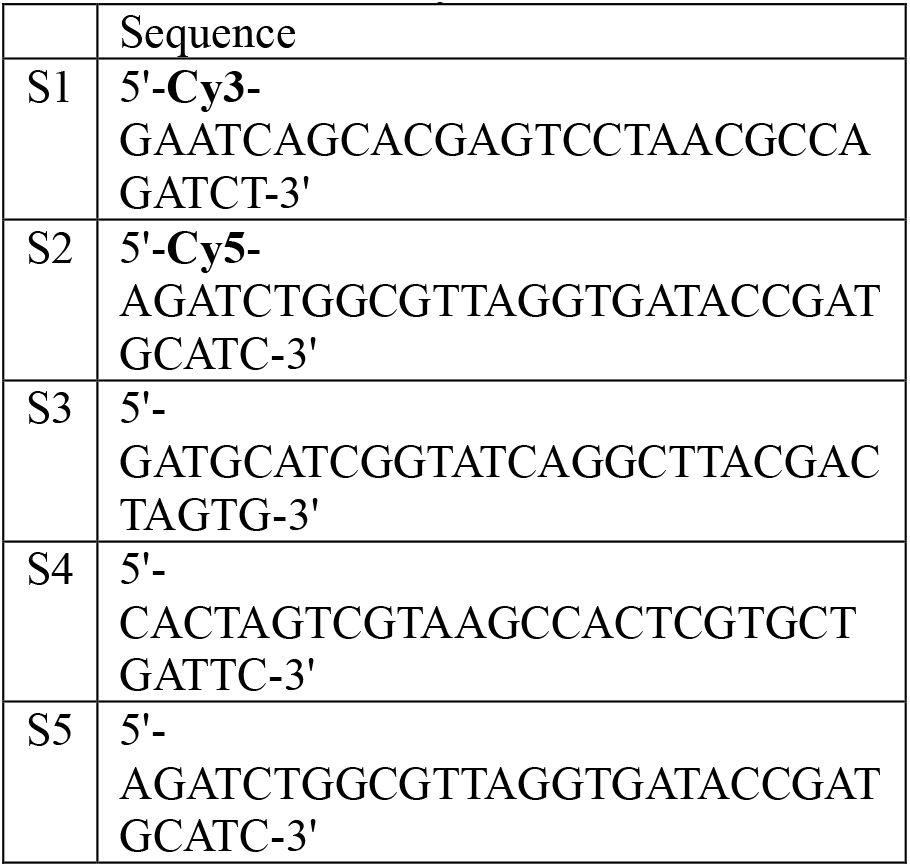
Oligonucleotide sequences used to assemble fluorescently labeled HJ J1.

### Purification of AOT

Commercially bought AOT [Bis(2-ethylhexyl) sulfosuccinate sodium salt] (Sigma-Aldrich) was purified using activated charcoal (Merck). AOT, pre-heated charcoal, and diethyl ether (Spectrochem) were mixed in a 300:100:1 ratio. The mixture was stirred using magnetic beads for 5-6 hours and filtered using a 0.2 μm syringe filter.^20^ This helps removing fluorescent impurities from AOT. AOT was dried using a vacuum desiccator by evaporating diethyl ether over several days. ^20^

### Preparation of RM

RM of W_0_ = 20 and 30 were prepared from a stock of AOT in isooctane (Spectrochem, UV grade) by mixing and vortexing the required amount of buffer containing HJs and MgCl_2_. The value of W_0_ is determined by the ratio of the added concentrations of the polar phase to AOT (W_0_ = [polar]/[AOT]). For Steadystate and time-resolved measurements, the final concentrations of AOT was15 mM, HJ was 10 nM, 20 mM Tris-HCl pH 7.5, and MgCl_2_ concentration was varied from 0.5 mM to 50 mM. Another set of HJ samples was prepared in buffer (20 mM Tris-HCl, pH 7.5) with varying MgCl_2_ concentration (0.5 mM to 50 mM).

### Steadystate fluorescence measurements

The steady-state fluorescence emission spectra were acquired using JASCO Spectrofluorometer FP-8500. Cy3 was excited by 532 nm laser and fluorescence emission was collected from 540 nm to 750 nm. FRET efficiency (E_FRET_) was calculated as E_FRET_ = I_A_/(I_D_+I_A_), where I_D_ are I_A_ the peak fluorescence intensities of the donor (Cy3) and acceptor (Cy5) fluorophores.

### Time-resolved data acquisition using TCSPC technique

Time-resolved fluorescence lifetime decays were obtained using TCSPC (FLS1000 Edinburgh Instruments) technique. Samples were excited with 510 nm, vertically polarised pulsed laser. The lifetime decays were collected at 565 nm at magic angle (54.8°) after passing through a long-pass filter with a cut-off of 550 nm. The instrument response function was measured as 300 ps using a scattering solution.

A general intensity decay *I*(*t*) can be fitted with a multi-exponential function like

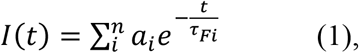

where *a*_*i*_ is the contribution of i^th^ species and τ_*Fi*_ is the fluorescence lifetime of the corresponding species. Earlier studies revealed a large conformational heterogeneity in HJ and a Gaussian distribution was required to perceive the isomer distribution.^7^ Thus, to incorporate isomer distribution over *n* number of species, Eq. 1 is modified as

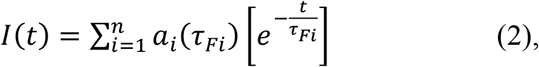

Where

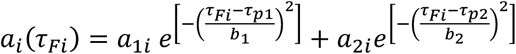

Here, *a*_1*i*_ and *a*_2 *i*_ are the relative contributions of the two distributions. τ_*p*1_ and τ_*p*2_ are the peak position of the Gaussian distribution, *b*_1_ and *b*_2_ are related to the width of the distribution. Average fluorescence lifetime (τ_*avg*_) was calculated as

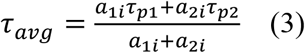

Here *a*_1*i*_ and *a*_2 *i*_ are taken as the peak intensity of the Gaussian distribution. FRET efficiency was calculated as

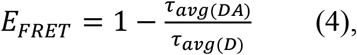

where τ_*avg(DA)*_is the average fluorescence lifetime of Cy3 in presence of Cy5 and τ_*avg(D)*_is that of the Cy3 in absence of Cy5.

## RESULTS AND DISCUSSIONS

### SteadyState FRET measurements shows higher FRET inside an RM

The ensemble-average FRET measurements of Cy3 and Cy5 labeled J1 were carried out at varying MgCl_2_ concentrations in aqueous buffer and inside RM of varying water pool size (Fig 1(B-F)). We used RM of W_0_ 20, and 30. Note that the water pool-size (diameter) of W_0_ = 20, and 30 are 7.2 nm and 11.1 nm respectively.^21^ The approximate size of the HJ used in our study fits well within W_0_ = 30, while W_0_ = 20 offers a much smaller space. In higher W_0_, e.g., 40 and 50, MgCl_2_ precipitated at higher concentrations. Thus, we limited our study to only W_0_ = 20, and 30 only. And finally, the FRET efficiencies (E_FRET_) are compared with HJ in buffer at all these ionic conditions(Fig 1(G)). In buffer, we observed that at 0.5 mM MgCl_2_ concentration, the E_FRET_ value is ∼ 0.15 and marginally increases to 0.2 as we systematically increased the MgCl_2_ concentration from 0.5 mM to 50 mM. At higher MgCl_2_ J1 is reported to preferentially populate *iso-I*, which is a high E_FRET_ population accordingly to our labelling strategy. As this is an ensemble-averaged measurement, we cannot ascertain the relative population of *iso-I* and *iso-II*.

However, we did observe a marginal increase in E_FRET_. Interestingly, when the HJ is put inside a confined environment of RM, E_FRET_ increases indicating that a restricted or constrained medium preferentially populates high FRET conformations of HJ (Fig 1(G)). The E_FRET_ increases 2.5 times for W_0_ = 20 and decreases as the water-pool size increases. E_FRET_ inside W_0_ = 30 matches with that for HJ in aqueous buffer indicating that the environment inside W_0_ = 30 is very similar to that of an unrestricted environment in aqueous buffer. Interestingly, for a particular environment, E_FRET_ remains almost the same for all the MgCl_2_ concentrations studied here (i.e., 0.5, 1.0, 5.0, 12.5 and 50 mM). We also observed that HJ J1 experiences a slightly more polar environment inside RM than that in a simple buffer. Most strikingly we observed that even at lower MgCl_2_ concentrations (0.5 mM and 1 mM, here) in a confined environment higher E_FRET_ is observed. We infer that in a confined environment, J1 prefers a conformation that is not an open X-like structure but a more compact one. On the contrary, in a diluted environment HJ is reported to adopt an open structure at lower salt concentration due to the electrostatic repulsion between arms. This observation may be relevant for understanding the recognition processes by junction binding proteins in confined cellular environment.

### Time-resolved measurements show lesser structural heterogeneity in the confined environment

Although it is widely accepted that the HJ undergoes a rapid transition between two stacked isomers through a very short-lived open structure, the structural heterogeneity during the transition has been shown through MD simulation.^22^ Additionally, time-resolved studies revealed a large conformational heterogeneity among the isomers.^7^ The lifetime of the donor (Cy3, here) is extremely sensitive to the distance between the donor-acceptor (D-A). Thus, if there is any distribution in D-A distance due to structural heterogeneity, that is expected to be reflected as a distribution of donor lifetime in the presence of an acceptor molecule. To confirm that, we have fitted the fluorescence intensity decay with two Gaussian distributions assuming there is *n* number of populations with peak fluorescence lifetimes of Cy3 as *τ*_*p1*_ and *τ*_*p2*_ (Eq. 2) (Fig. 2(A-C)). J1 exhibited two distinct lifetime distributions, a faster one, τ_*p*1_ and a slower one, τ_*p*2_ (Table 2). According to our labeling scheme, faster lifetime may be associated with *iso-I* (high-FRET) population and the longer lifetime can be associated with a mixed population of X-like open conformation and *iso-II* (low-FRET). The change in E_FRET_ with MgCl_2_ is shown in Fig 2(D). It is to be noted that the maximum E_FRET_ calculated here is much lesser than the estimated one (Fig 1(A). This can be the averaging effect between two isomers. Like our observation in steadystate measurements, we observed that even at lower salt concentrations, the E_FRET_ increases almost 2 times when the HJ is placed inside a RM. Additionally, E_FRET_ in buffer increases with increasing salt concentration. However, once the HJ is placed within the RM, E_FRET_ only marginally changes upon increasing salt concentration. This might signify that once placed within a confined environment, the effect of ionic conditions is a lesser governing factor than confinement. This also corroborates the presence of a higher FRET population even at lower MgCl_2_ concentrations inside the confined environment of RM. Fig. 3(A-E) shows the Gaussian distribution of the lifetimes at different MgCl_2_ concentrations in different environments. We observed that the width of the Gaussian distribution related to τ_*p*1_ is much narrower than that of τ_*p*2_ (Fig 3(F)). This is expected as the distribution of τ_*p*2_ is considered to have population for both *iso-II* and open X-like conformation. We also observed a higher relative amplitude of τ_*p*1_ compared to τ_*p*2_ . This corroborates with previous studies reporting a higher population of *iso-I* for J1.^7,14^ We observed that overall, the width of the distribution related to τ_*p*2_ is much higher in buffer compared to τ_*p*1_ and it decreases significantly when the HJ is confined inside an RM. This might be due to confinement effect. As HJ experience a restricted environment inside a RM, the conformational heterogeneity is also restricted. Although we observed both the populations in all three environments, the relative amplitude of slower lifetime increases in confined environment with an overall increase in the E_FRET_ compared to buffer. We infer that in a confined environment, HJ prefers to stay in stacked conformation and the effect is more prominent at lower MgCl_2_ concentration.

**Table 2.** Fitted parameters of the time-resolved decays.

| | [MgCl<sub>2</sub>]<br>(mM) | Only D (Cy3) | | | | | D-A (Cy3-Cy5) | | | | | $E_{FRET}$ |
| --- | --- | --- | --- | --- | --- | --- | --- | --- | --- | --- | --- | --- |
| | | $a_1$<br>(peak) | $\tau_{p1}$<br>(ns) | $a_2$<br>(peak) | $\tau_{p2}$<br>(ns) | $\tau_{avg(D)}^*$<br>(ns) | $a_1$<br>(peak) | $\tau_{p1}$<br>(ns) | $a_2$<br>(peak) | $\tau_{p2}$<br>(ns) | $\tau_{avg(DA)}^*$<br>(ns) | |
| **W<sub>0</sub> = 20** | 0.5 | 0.58 | 1.4 | 0.12 | 2.1 | 1.5 | 0.38 | 0.6 | 0.22 | 1.5 | 1.0 | 0.36 |
|  | 1.0 | 0.62 | 1.4 | 0.11 | 2.1 | 1.5 | 0.34 | 0.6 | 0.19 | 1.5 | 0.9 | 0.40 |
|  | 5.0 | 0.61 | 1.4 | 0.07 | 2.1 | 1.4 | 0.38 | 0.6 | 0.20 | 1.5 | 0.9 | 0.38 |
|  | 12.5 | 0.64 | 1.3 | 0.10 | 2.1 | 1.4 | 0.46 | 0.6 | 0.22 | 1.5 | 0.9 | 0.40 |
|  | 50.0 | 0.67 | 1.2 | 0.03 | 2.9 | 1.3 | 0.35 | 0.5 | 0.18 | 1.4 | 0.8 | 0.41 |
| **W<sub>0</sub> = 30** | 0.5 | 0.61 | 1.3 | 0.12 | 2.4 | 1.5 | 0.44 | 0.7 | 0.20 | 1.5 | 0.9 | 0.36 |
|  | 1.0 | 0.82 | 1.4 | 0.08 | 2.3 | 1.4 | 0.37 | 0.7 | 0.18 | 1.4 | 0.9 | 0.37 |
|  | 5.0 | 0.86 | 1.4 | 0.06 | 2.9 | 1.5 | 0.33 | 0.6 | 0.20 | 1.5 | 1.0 | 0.35 |
|  | 12.5 | 0.81 | 1.3 | 0.08 | 2.2 | 1.4 | 0.42 | 0.7 | 0.17 | 1.5 | 0.9 | 0.38 |
|  | 50.0 | 0.69 | 1.0 | 0.16 | 2.0 | 1.2 | 0.50 | 0.6 | 0.17 | 1.4 | 0.8 | 0.36 |
| **Buffer** | 0.5 | 0.42 | 0.8 | 0.19 | 2.2 | 1.2 | 0.35 | 0.6 | 0.14 | 1.9 | 1.0 | 0.20 |
|  | 1.0 | 0.46 | 0.8 | 0.20 | 2.3 | 1.2 | 0.39 | 0.6 | 0.13 | 1.9 | 1.0 | 0.23 |
|  | 5.0 | 0.37 | 0.7 | 0.05 | 2.2 | 0.9 | 0.39 | 0.5 | 0.08 | 1.6 | 0.7 | 0.19 |
|  | 12.5 | 0.83 | 0.7 | 0.02 | 2.1 | 0.7 | 0.40 | 0.5 | 0.07 | 1.2 | 0.6 | 0.23 |
|  | 50.0 | 0.80 | 0.7 | 0.03 | 2.0 | 0.7 | 0.36 | 0.4 | 0.09 | 1.0 | 0.5 | 0.31 |

**Fig. 2.**
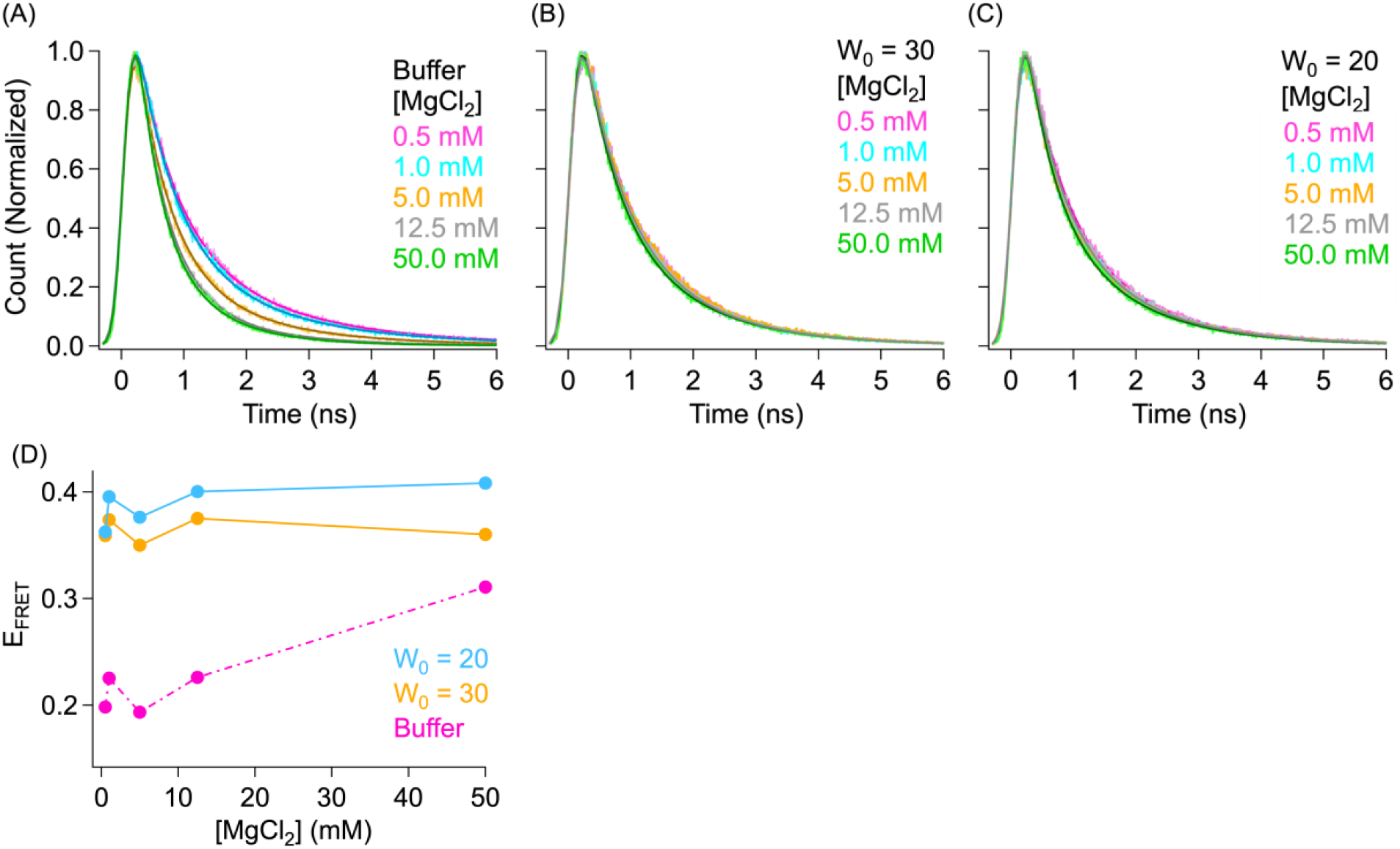
Time-resolved fluorescence decay of Cy3-Cy5 labeled J1 in (A) buffer and inside RMs (B) W_0_ = 30 and (C) W_0_ = 20 with 0.5 mM, 1.0 mM, 5 mM, 12.5 mM and 50 mM MgCl_2_. (D) Change in FRET efficiency with MgCl_2_ in different environment. The fits with Gaussian distribution model (Eqn. 2) are shown in darker shades.

**Fig. 3.**
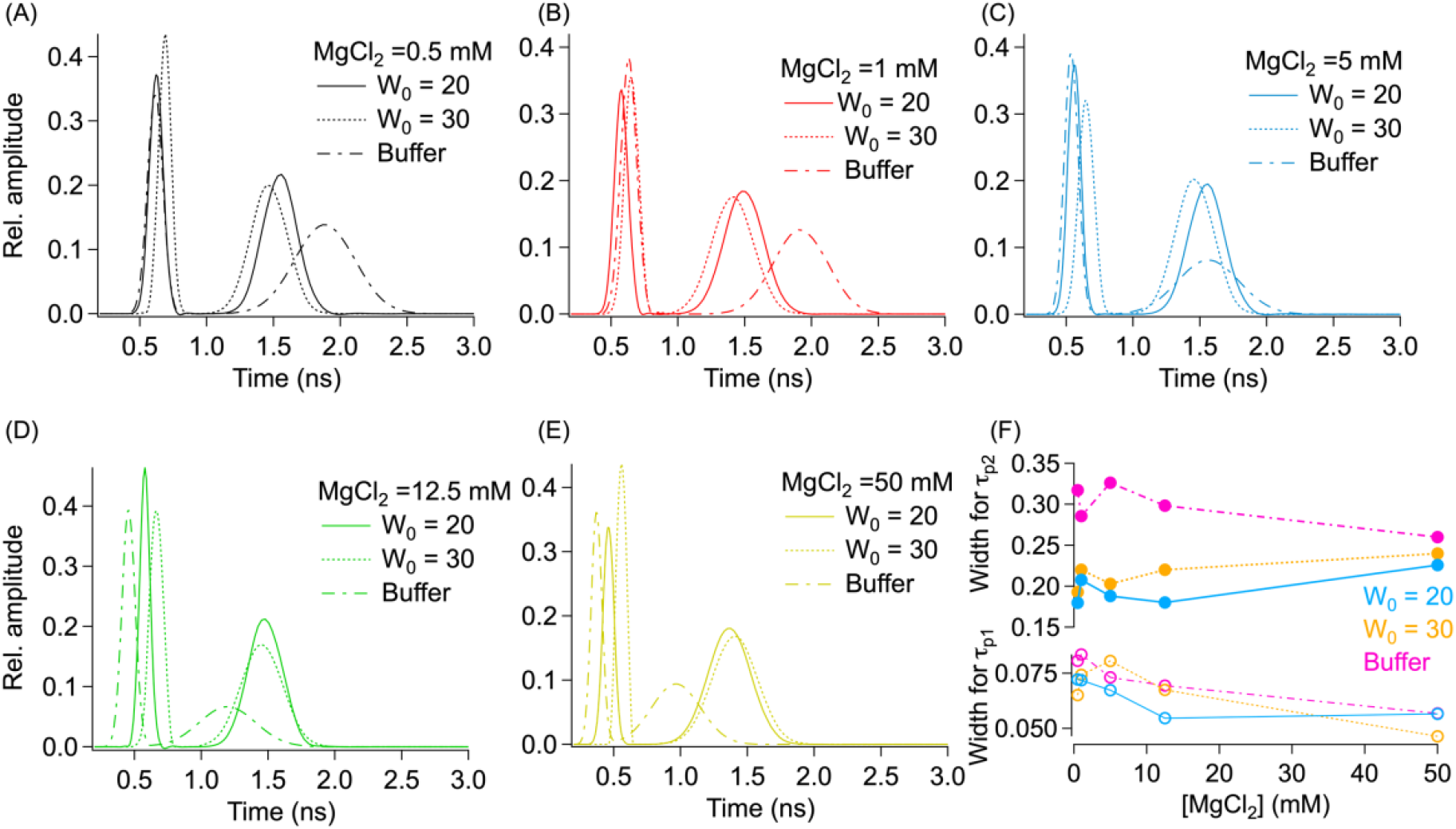
Gaussian distribution of the lifetime of Cy3-Cy5 labeled J1 in buffer and inside RMs with (A) 0.5 mM, (B) 1.0 mM, (C) 5 mM, (D) 12.5 mM and (E) 50 mM MgCl_2_. (F) Change width of the Gaussian distribution with MgCl_2_.

## CONCLUSION

The role of confined/restricted environment on the conformation of biomolecules is well established. However, the insight on fourway HJ in a confined environment is incomplete. Numerous studies using synthetic HJ have shown that it adopts two global conformers, open X-like and stacked structures. At lower ionic condition, the inter-arm electrostatic repulsion directs the HJ to adopt an extended open X-like structure. The presence of an adequate amount of divalent metal ions minimizes the electrostatic repulsion and HJ forms more compact stacked structures. The stacked conformers can exist in two isomeric forms and are distinguished by which duplex arms are stacked on each other. While the isomer transition and resolution of the HJ by enzymes have been well investigated, there is a dearth of knowledge about the behavior of HJ in a confined intracellular environment.

We employed fluorescence-based techniques to investigate the behavior of HJ inside RM which offers an excellent alternative for confined space. Compared to HJ in aqueous buffer, our study shows that in a confined environment, HJ prefers to adopt stacked conformer. The time-resolved fluorescence data shows a decrease in heterogeneity in the lifetime of the donor in presence of the acceptor which is directly reflective of the inter-arm distance. More interestingly, we observed that inside the confined environment of RM, HJ populates stacked structures even at low MgCl_2_ concentrations. Such confinement-induced changes in conformer distribution can be expected to affect the activity of the junction binding proteins. Overall, our results provide an insight into the conformer distribution of HJ in a confined space.

## ACKNOWLEDGMENTS

The work has been supported by IISER Tirupati. P. A. and S. H. K thank IISER Tirupati for the fellowship. We thank Dr. Janardan Kundu and Dr. Jatish Kumar for kindly allowing us to access the time-resolved set-up.

## Notes

The authors declare no competing interest.

## REFERENCE

(1) Liu, Y. W. C. S. Nat Rev Mol Cell Biol 2004, 5, 933–944.

(2) Wright, W. D.; Shah, S. S.; Heyer, W. D. J Biol Chem 2018, 293, 10524–10535.

(3) Eichman, B. F.; Vargason, J. M.; M Mooers, B. H.; Shing Ho, P. Proc Natl Acad Sci USA 2000, 97, 3971–3976.

(4) Ray, S.; Pal, N.; Walter, N. G. Nucleic Acids Res 2021, 49, 2803–2815.

(5) McKinney, S. A.; Déclais, A. C.; Lilley, D. M. J.; Ha, T. Nat Struct Biol 2003, 10, 93–97.

(6) Joo, C.; McKinney, S. A.; Lilley, D. M. J.; Ha, T. J Mol Biol 2004, 341, 739–751.

(7) Miick, S. M.; Fee, R. S.; Millar, D. P.; Chazin, W. J.; Zimm, B. H. Proc Natl Acad Sci USA 1997; 94, 9080–9084.

(8) Zhou, R.; Yang, O.; Déclais, A. C.; Jin, H.; Gwon, G. H.; Freeman, A. D. J.; Cho, Y.; Lilley, D. M. J.; Ha, T. Nat Chem Biol 2019, 15, 269–275.

(9) Khuu, P. A.; Voth, A. R.; Hays, F. A.; Ho, P. S. J Mol Recognit 2006, 19, 234–242.

(10) Mahmoud, R.; Dhakal, S. J Phys Chem B 2022, 126, 7252–7261.

(11) Zhang, F.; Simmons, C. R.; Gates, J.; Liu, Y.; Yan, H. Angewandte Chemie - International Edition 2018, 57, 12504–12507.

(12) Simmons, C. R.; Zhang, F.; MacCulloch, T.; Fahmi, N.; Stephanopoulos, N.; Liu, Y.; Seeman, N. C.; Yan, H. J Am Chem Soc 2017, 139, 11254–11260.

(13) Zheng, J.; Birktoft, J. J.; Chen, Y.; Wang, T.; Sha, R.; Constantinou, P. E.; Ginell, S. L.; Mao, C.; Seeman, N. C. Nature 2009, 461, 74–77.

(14) Gibbs, D. R.; Dhakal, S. Biochemistry 2018, 57, 3616–3624.

(15) Sung, H. L.; Nesbitt, D. J. J Phys Chem B 2021, 125, 13147–13157.

(16) Cino, E. A.; Karttunen, M.; Choy, W. Y. PLoS One 2012, 7, e49876.

(17) Van Horn, W. D.; Ogilvie, M. E.; Flynn, P. F. J Am Chem Soc 2009, 131, 8030–8039.

(18) Khamari, L.; Mukherjee, S. J Phys Chem Letts 2022, 13, 8169–8176.

(19) Ho, M. C.; Chang, C. W. RSC Adv 2014, 4, 20531–20534.

(20) Pal, N.; Verma, S. D.; Singh, M. K.; Sen, S. Anal Chem 2011, 83, 7736–7744.

(21) Pal, N.; Verma, S. D.; Singh, M. K.; Sen, S. Anal Chem 2011, 83, 7736–7744.

(22) Yu, J.; Ha, T.; Schulten, K. Nucleic Acids Res 2004, 32, 6683–6695.

